# Cuticular Hydrocarbons on Old Museum Specimens of the Spiny Mason Wasp, *Odynerus spinipes*, (Hymenoptera: Vespidae: Eumeninae) Shed Light on the Distribution and on Regional Frequencies of Distinct Chemotypes

**DOI:** 10.1101/2020.09.05.283291

**Authors:** Victoria C. Moris, Katharina Christmann, Aline Wirtgen, Sergey A. Belokobylskij, Alexander Berg, Wolf-Harald Liebig, Villu Soon, Hannes Baur, Thomas Schmitt, Oliver Niehuis

## Abstract

The mason wasp *Odynerus spinipes* shows an exceptional case of intrasexual CHC dimorphism. Females of this species express one of two cuticular hydrocarbon (CHC) profiles (chemotypes) that differ qualitatively and quantitatively from each other. The ratio of the two chemotypes was previously shown to be close to 1:1 at three sites in Southern Germany, which might not be representative given the Palearctic distribution of the species. To infer the frequency of the two chemotypes across the entire distributional range of the species, we analyzed with GC-MS the CHC profiles of 1,042 dry-mounted specimens stored in private and museum collections. We complemented our sampling by including 324 samples collected and preserved specifically for studying their CHCs. We were capable of reliably identifying the chemotypes in 91% of dry-mounted samples, some of which collected almost 200 years ago. We found both chemotypes to occur in the Far East, the presumed glacial refuge of the species, and their frequency to differ considerably between sites and geographic regions. The geographic structure in the chemotype frequencies could be the result of differential selection regimes and/or different dispersal routes during the colonization of the Western Palearctic. The presented data pave the route for disentangling these factors by providing information where to geographically sample *O. spinipes* for population genetic analyses. They also form the much-needed basis for future studies aiming to understand the evolutionary and geographic origin as well as the genetics of the astounding CHC dimorphism that *O. spinipes* females exhibit.

## Introduction

The cuticular hydrocarbon (CHC) profiles of insects are generally thought to be species-specific (Bagnères and Wicker-Thomas 2010), with profile differences between species often being both qualitative and quantitative. Intraspecific CHC profile differences are most often observed between sexes and between samples from different populations, but CHC profiles are also known to change with the insect’s age, mating and fertility status, and — in case of eusocial insects — with the individuals caste, task, and colony membership (Sledge et al. 2001; Cuvillier-Hot et al. 2001; Greene and Gordon 2003; Hugo et al. 2006; Jackson et al. 2007; Ichinose and Lenoir 2009; Martin and Drijfhout 2009a; Nunes et al. 2009; Blomquist and Bagnères 2010; Kuo et al. 2012; Polidori et al. 2017; Korb 2018). While intersexual CHC dimorphism is common in insects, intrasexual CHC dimorphism has been found in only a few species, one of which is the spiny mason wasp, *Odynerus spinipes* (Linnaeus, 1758) (Strohm et al. 2008a, b; Marten et al. 2009; Martin et al. 2010; de Oliveira et al. 2011; Wurdack et al. 2015).

The mason wasp *O. spinipes* is one of comparatively few known solitary insects that show conspicuous intrasexual dimorphism: females are able to express one of two CHC profiles (also referred to as chemotypes) that differ in more than 70 chemical compounds from each other (Wurdack et al. 2015). These qualitative differences are primarily due to the presence or absence of alkenes with double bonds at specific positions: chemotype 1 is characterized by alkenes with double bonds at positions 5, 7, and 9 of the hydrocarbon chain (hereinafter referred to as *udp* alkenes), whereas chemotype 2 is characterized by alkenes with double bonds at the positions 8, 10, 12, and 14 of the hydrocarbon chain (hereinafter referred to as *edp* alkenes) (Wurdack et al. 2015). No CHC profile has been recorded so far that was interpreted as intermediate between chemotype 1 and chemotype 2. *O. spinipes* males only express one chemotype, which show only minor differences to chemotype 1 (Wurdack et al. 2015).

Wurdack et al. (2015) suggested that the CHC dimorphism of *O. spinipes* females is the result of an evolutionary arms race between the spiny mason wasp and two kleptoparasitic cuckoo wasps: *Pseudochrysis neglecta* (Shuckard 1837) (formerly in the genus *Pseudspinolia*; Rosa et al. 2017) and *Chrysis mediata* (Linsenmaier 1951) (Hymenoptera: Chrysididae). Each of the two kleptoparasites seem to chemically mimic one of the two chemotypes. As the two cuckoo wasps are distantly related to each other (Pauli et al. 2019), they likely evolved their exploitation of *O. spinipes* as host independently from each other. Because Wurdack et al. (2015) found females expressing the two chemotypes to occur sympatrically and observed a balanced (roughly 1:1) frequency of the two chemotypes at all three *O. spinipes* nesting sites that they studied in Southern Germany, the authors hypothesized that the genetic information for expressing the two chemotypes is maintained in *O. spinipes* populations by balancing selection (Wurdack et al. 2015). Given the vast distributional range of *O. spinipes*, occurring from Spain in the West to the Pacific coast in the Far East (Gusenleitner 1998; Woydak 2006; present study), the chemotype frequencies determined by Wurdack et al. (2015) are not necessarily representative for the majority of populations of this species, though. Representatively sampling populations of a species with a huge distributional range is often difficult to achieve, however, due to the associated costs for traveling. A possible solution of this problem could be the exploitation of samples deposited in museum collections.

In this study, we (1) test whether or not dry-mounted samples of *O. spinipes* collected by entomologists during the last 200 years and stored in private and museum collections carry a sufficient quantity of CHCs to determine their chemotypes. For this purpose, we analyzed CHC extracts of 1,042 dry-mounted *O. spinipes* samples using gas-chromatography coupled with mass-spectrometry (GC-MS). Since CHCs are — due to of their high molecular mass — poor volatiles and seem to be chemically stable over time (Page et al. 1990; Martin et al. 2009b), museum specimens could indeed proof to be a valuable resource for studying CHC profiles in a biogeographic context. We complemented the data set with 324 freeze-kill fresh samples collected and preserved specifically for studying their CHCs and further exploited the collected data to (2) test whether or not the frequency of the two chemotypes that *O. spinipes* females are able to express is roughly 1:1 in populations from the entire distributional range of the species. Since our sampling also includes male specimens, we furthermore (3) test whether or not *O. spinipes* males express consistently only one chemotype. The collected data finally allowed us to assess more thoroughly than Wurdack et al. (2015) (4) whether or not *O. spinipes* females that exist are able to express a CHC profile that is intermediate between that of chemotype 1 and that of chemotype 2. The reported data form the much-needed basis for studies that seek to understand the evolutionary and geographic origin as well as the genetics of the astounding qualitative CHC dimorphism that the spiny mason wasp exhibits.

## Methods and Materials

### Samples

We studied 1,042 dry-mounted specimens stored in the following private or public collections: Deutsches Entomologisches Institut (Eberswalde, Germany; 74 wasps), Museum für Naturkunde (Berlin, Germany, 159 wasps), Naturhistorisches Museum (Wien, Austria; 47 wasps), private collection O. Niehuis (Freiburg, Germany; 173 wasps), Biozentrum of the Oberösterreichisches Landesmuseum (Linz, Austria; 454 wasps), private collection C. Schmid-Egger (Berlin, Germany; 27 wasps), Senckenberg Naturmuseum (Frankfurt a. M., Germany; 44 wasps), Senckenberg Museum für Naturkunde (Görlitz, Germany; 20 wasps), Staatliches Museum für Naturkunde (Stuttgart, Germany; 49 wasps), Zoological Institute of the Russian Academy of Sciences (St. Petersburg, Russia; 52 wasps), Institute of Biology and Biomedicine at the Lobachevsky State University (Nizhny Novgorod, Russia; 2 wasps), and Zoologische Staatssammlung (Munich, Germany; 97 wasps). We additionally studied 324 samples collected and directly frozen at -20 °C by us at locations in Belgium (105 wasps), Sweden (26 wasps), Estonia (26 wasps), and Southern Germany (167 wasps). More information about the samples analyzed, including sampling sites and associated geocoordinates, are provided in the Supplementary Table S1. Note that in those instances in which the label information of samples did not include the geocoordinates of the sampling site, we inferred this information using GoogleMaps (Google, Mountain View, CA, USA). We analyzed 1,366 specimens in total (i.e., dry-mounted and fresh ones), of which 1,009 were females and 357 were males.

### Cuticular Hydrocarbon Extraction

Females preserved in insect boxes were individually immersed in n-hexane (SupraSolv n-hexane for gas chromatography, Merck KGaA, Germany, or Rotipuran n-hexane, Carl Roth GmbH, Karlsruhe, Germany) for 2 or for 10 minutes, depending on the specimen (after analyzing the first batch of dry-mounted specimens, we increased the extraction time from 2 to 10 minutes to increase the CHC yield). All CHC extracts were subsequently reduced to 75 µL by evaporating them under a gentle constant flow of nitrogen. The CHC extracts were stored at -20 °C before proceeding with their chemical analysis. The CHCs of all female wasps collected by us in Belgium (Tenneville) or in Germany (Büchelberg) were either extracted for 10 minutes with n-hexane or were sampled with a Solid-Phase Micro Extraction (SPME) fiber (Supelco, coating: polydimethylsiloxane, 100 µm, Sigma Aldrich, Bellefonte, PA, USA). Females sampled with SPME fibers were anesthetized either by exposing them for 1 minute to CO_2_ (females sampled in 2016) or by cooling them for 3 minutes down at -20 °C (females sampled in 2017 and 2018). Conditioned SPME fibers were scrubbed for 2 minutes against the wasps’ metasoma.

### Chemical Analysis

GC-MS analyses were conducted with a 7890B gas chromatograph system (Agilent Technologies, Santa Clara, CA, USA) equipped with a DB-5 column (30 m × 0.25 mm ID, df = 0.25 μm, J & W Scientific, Folsom, USA) coupled with a 5977B mass selective detector (Agilent Technologies, Santa Clara, CA, USA), using helium at a constant flow of 1 ml min^-1^ as a carrier gas. The CHC extracts of ten wasps collected in Tenneville (Belgium) were analyzed with a gas chromatograph-flame ionization detector (Shimadzu GC-2010 system) equipped with a SLB-5 ms non-polar capillary column (5% phenyl (methyl) polysiloxane stationary phase; 30 m column length; 0.25 mm inner diameter; 0.25 μm film thickness). We applied in both instruments the following temperature program: start temperature 40 °C, increased by 10 °C per minute up to 300 °C, which was maintained for 10 min. The injection port was set at 250 °C and operated in spitless mode for 1 minute. The electron ionization mass spectra were acquired at an ionization voltage of 70 eV (source temperature: 230 °C). Total intensity chromatograms and mass spectra were inferred with the software MSD Enhanced ChemStation F.01.03.2357 for Windows (Agilent Technologies, Böblingen, Germany). Individual cuticular hydrocarbons were identified based on their diagnostic ions. To control the sensitivity of the GC-MS, we injected every week and before running a batch of samples a C7–C40 saturated alkanes standard (Sigma-Aldrich, Steinheim, Germany) into the GC-MS.

The two chemotypes that *O. spinipes* females are known to express were identified by studying the presence/absence of diagnostic alkenes, each of which with a specific retention time, in the acquired chromatograms. According to Wurdack et al. (2015), chemotype 1 is characterized by alkenes with double bonds at uneven positions of the hydrocarbon chain (*udp* alkenes), whereas chemotype 2 is characterized by alkenes with double bonds at even positions of the hydrocarbon chain (*edp* alkenes). Note that it is easily possible to distinguish the two chemotypes by carefully studying total ion chromatograms and mass spectra, even without derivatization of the CHC extracts: alkenes of chemotype 1 appear at slightly later retention times than the corresponding alkenes of chemotype 2. The difference is particularly prominent when focusing on the most abundant alkenes on the cuticle of *O. spinipes* females (i.e., those with hydrocarbon chain lengths of 25, 27, and 29; Fig. 1). We nonetheless checked the double bond position of diagnostic alkenes in a subset of samples by sample derivatization applying a custom protocol derived from that given by Dunkelblum et al. 1985 and Carlson et al. 1989. Specifically, we inferred the position of double bonds on the cuticle of 146 dry-mounted *O. spinipes* females originating from collections and of 21 *O. spinipes* females freshly collected by us in the field (Table S3). For this purpose, we first concentrated all CHC extract volumes to 120 µL each and then mixed each of them with dimethyl disulfide in a 1:1 ratio. We added to the resulting volume (240 µL) 85 µL of a 5% iodine solution in diethyl ether. The reaction volume was thoroughly mixed by hand-shaking it and then kept for 12–24 h at 60 °C. We subsequently added as many drops of a 5% sodium thiosulfate solution in water until the solution turned transparent. Note that the solution was thoroughly shaken by hand after each added drop. Finally, we isolated the organic (upper) phase, transferred it to a new vial, concentrated it under a gentle constant flow of nitrogen, and transferred it into an insert (Agilent Technologies Inc., Santa Clara, CA, U.S.A.; 25 ml glass with polymer feet). The derivatized CHC extracts were analyzed with the 7890B gas chromatograph system and applying the same instrument settings as described above.

**Fig. 1.**
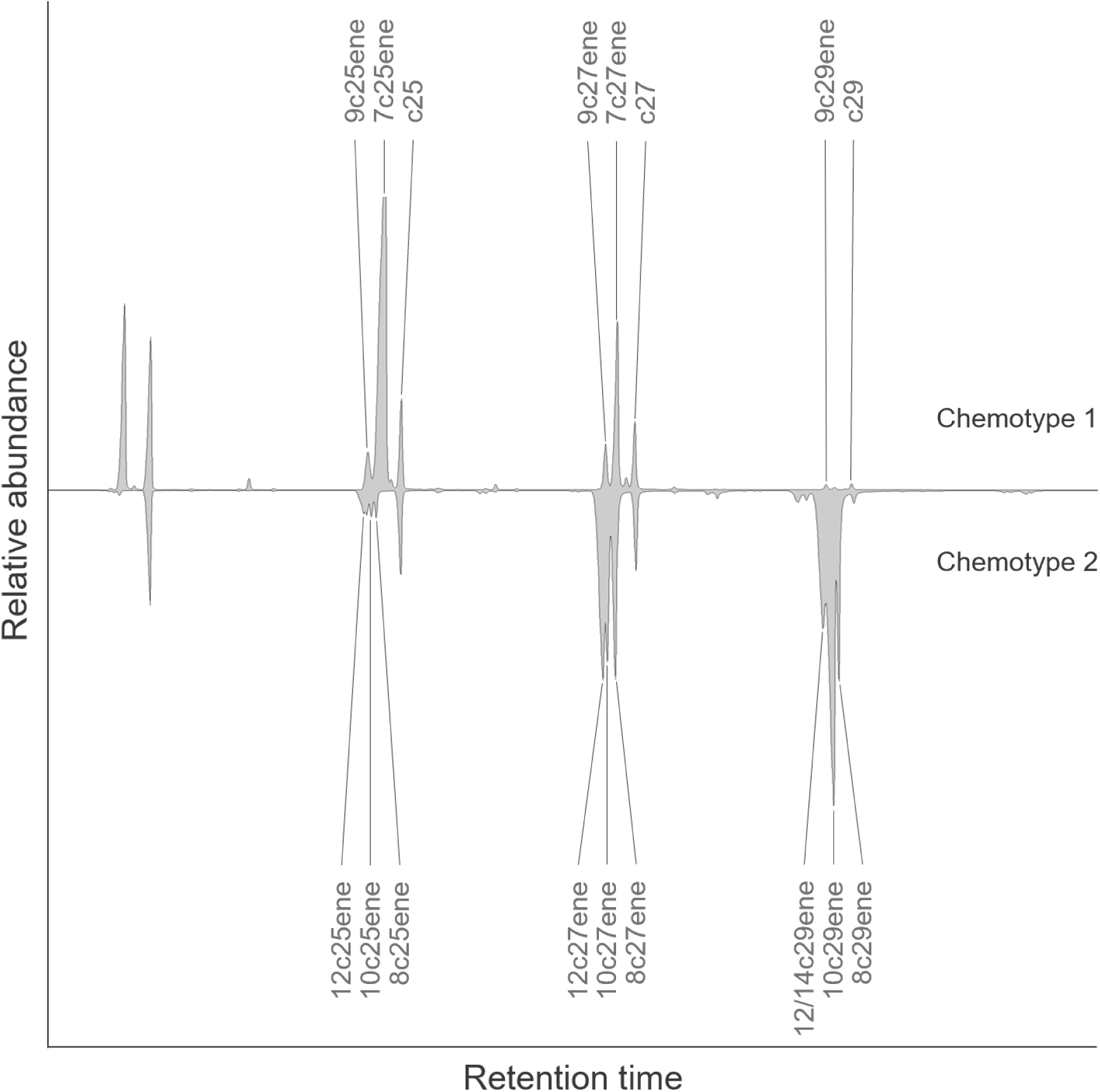
Chromatograms of frozen sampled CHC extracts of *Odynerus spinipes* females stored at - 20 °C. The retention time differ slightly between alkenes of females expressing chemotype 1 (top), which have the double bond at an uneven position, compared to alkenes of females expressing chemotype 2 (bottom), which have the double bond at an even position.

To determine whether or not the quality of CHC extracts from dry-mounted specimens depended on the number of years between the collection of the specimen and its CHC extraction, we performed a logistic regression with the software R version 3.4.1 (R Core Team 2017), using the function glm (Dobson 1990) of the package stats version 3.6.2.

### Maps and Graphics

Records of *O. spinipes* females in close spatial proximity to each other were grouped to increase the sample size within a group prior to calculating chemotype frequencies (detailed in the Supplementary Table S1). Specifically, females collected in Central Europe were grouped so that the number of samples under a circle area with 125 km diameter (covering a surface area of 12,272 km^2^) was maximized (Fig. S1). Because regional sample sizes outside Central Europe were typically small, we chose a circle diameter of 420 km for grouping females collected outside Central Europe. The procedure resulted in a total of 53 groups, 27 of which located in Central Europe (Table S2). To visualize the chemotype frequencies in each of the 53 groups, we plotted pie charts and mapped them onto a map of Central Europe and onto a map of the Palearctic region, with the size of the pie charts proportional to number of female samples included in the respective group. The center of the pie charts is positioned at the location in the circle with the shortest distance to all other locations in the circle. The diameter of the pie charts is the log_10_ of the number of females in the group. While groups in Central Europe always contained more than two samples, group sizes in the remaining parts of the Palearctic contained in some instances only one sample. To nonetheless be able to visualize these records on the Palearctic map, we added the value 0.5 to all log_10_ values. The pie chart diameter of groups composed only of one sample was consequently 0.5 and not zero. Note that we divided the log_10_ values by 4 when plotting pie charts onto the map of Central Europe to reduce the pie charts’ diameter. All maps were drawn with the software R version 3.4.1 (R Core Team 2017), using the package “maps” version 3.3.0. Pie charts were plotted onto the maps with the aid of the R package “mapplots” version 1.5.1 (Gerritsen 2013).

### Morphometric Analysis

Since we found the CHC profiles of most dry-mounted *O. spinipes* females likely expressing chemotype 2 to differ in their CHC composition for the typical chemotype 2 CHC composition described by Wurdack et al. (2015), we conduced morphometric analyses on female samples to assess the possibility that our sampling included a cryptic species. For this purpose, we studied 223 dry-mounted *O. spinipes* females from five different localities, including those at which females with deviant CHC profiles were recorded. Specifically, we analyzed (A) 28 females from the vicinity of Frankfurt a. M. (Germany; 7 expressing chemotype 1, 21 expressing the deviant chemotype 2), (B) 32 females from the vicinity of Leipzig (Germany; 9 expressing chemotype 1, 23 expressing the deviant chemotype 2), (C) 99 females collected in South Bohemia (Czech Republic;64 expressing chemotype 1, 7 expressing chemotype 2, 28 expressing the deviant chemotype 2), (D) 50 females from the vicinity of Syrovice (Czech Republic; 2 expressing chemotype 1, 48 expressing the deviant chemotype 2), and (E) 14 females collected in the Far East (Russia; all expressing the deviant chemotype 2). We followed the protocol for morphometric analysis given by Wurdack et al. (2015) with slight modifications; specifically, we selected and considered in the final analysis 15 morphological characters from the 16 studied characters based on a reliability test of the characters (see below; Table 1, Fig. S2). We followed the terminology for naming morphological structures given by Gibson (<u>1997</u>). A list of all quantified morphometric characters is given in Table 1.

**Table 1.**
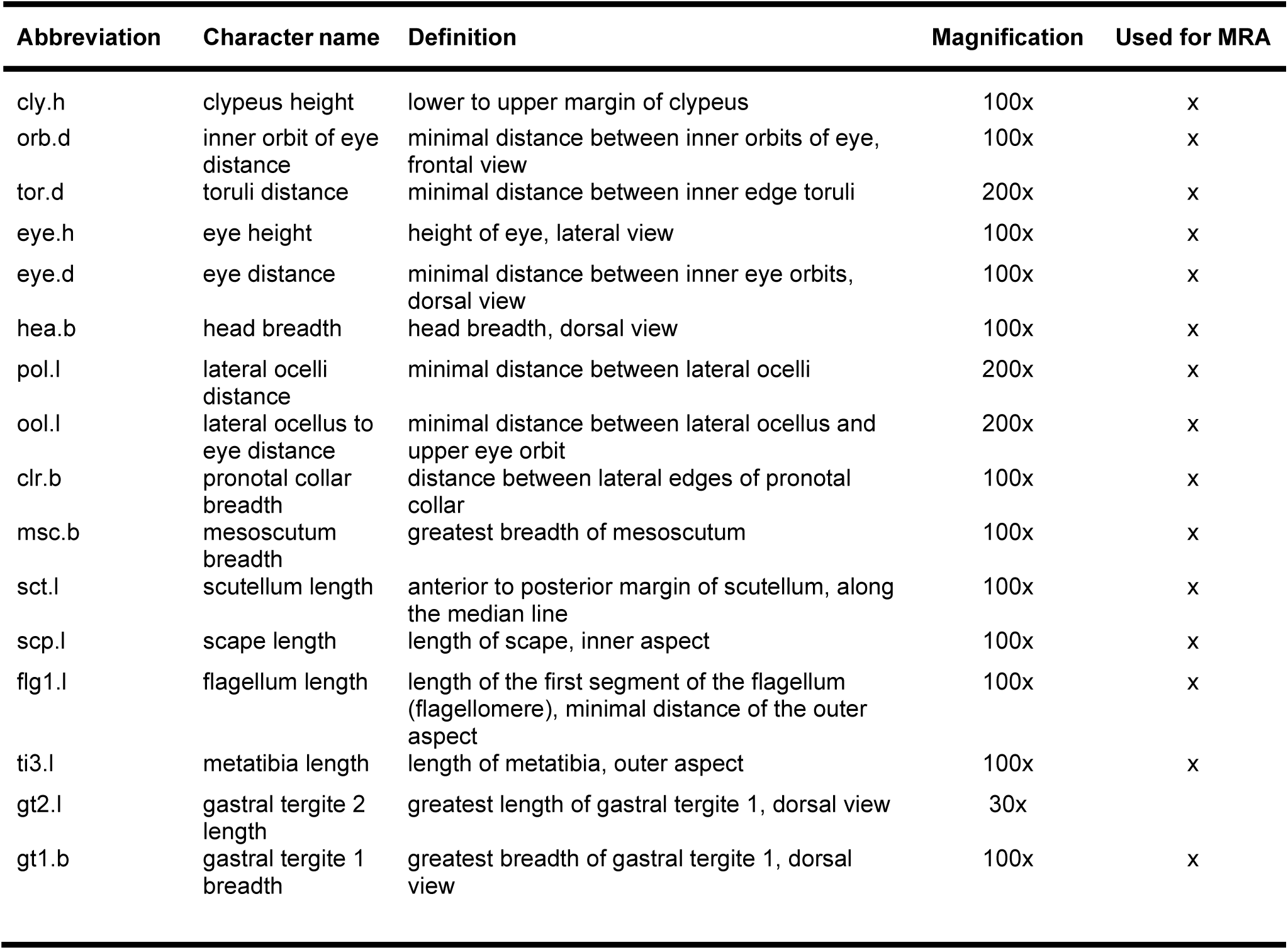
Abbreviations, names, definitions, and magnifications used to morphometrically study 16 characters of *Odynerus spinipes* females, and those used in the multivariate ratio analysis (MRA)

| Abbreviation | Character name | Definition | Magnification | Used for MRA |
| --- | --- | --- | --- | --- |
| cly.h | clypeus height | lower to upper margin of clypeus | 100x | x |
| orb.d | inner orbit of eye distance | minimal distance between inner orbits of eye, frontal view | 100x | x |
| tor.d | toruli distance | minimal distance between inner edge toruli | 200x | x |
| eye.h | eye height | height of eye, lateral view | 100x | x |
| eye.d | eye distance | minimal distance between inner eye orbits, dorsal view | 100x | x |
| hea.b | head breadth | head breadth, dorsal view | 100x | x |
| pol.l | lateral ocelli distance | minimal distance between lateral ocelli | 200x | x |
| ool.l | lateral ocellus to eye distance | minimal distance between lateral ocellus and upper eye orbit | 200x | x |
| clr.b | pronotal collar breadth | distance between lateral edges of pronotal collar | 100x | x |
| msc.b | mesoscutum breadth | greatest breadth of mesoscutum | 100x | x |
| sct.l | scutellum length | anterior to posterior margin of scutellum, along the median line | 100x | x |
| scp.l | scape length | length of scape, inner aspect | 100x | x |
| flg1.l | flagellum length | length of the first segment of the flagellum (flagellomere), minimal distance of the outer aspect | 100x | x |
| ti3.l | metatibia length | length of metatibia, outer aspect | 100x | x |
| gt2.l | gastral tergite 2 length | greatest length of gastral tergite 1, dorsal view | 30x |  |
| gt1.b | gastral tergite 1 breadth | greatest breadth of gastral tergite 1, dorsal view | 100x | x |

Each character was photographed with a Keyence VHX 2000 digital photo-microscope and a VH-Z20R/W zoom lens (Keyence, Japan, USA) at different magnifications (30x, 100x, 200x) depending on the photographed character. To ensure that the magnification did not change between photographs showing the same character in different samples, we conducted all measurements requiring the same magnification in one session without altering the magnification in-between and used an eye-piece micrometer (12 mm subdivided into 120 units; 1,000 µm corresponded to 888 pixels) to calibrate photographs. To remove additional variation possibly caused by fluctuating asymmetry (Palmer and Strobeck 1986; Bechshøft et al. 2008), we measured characters on the left-hand side, when it was possible. Additionally, we processed the samples in random order to avoid the possibility of systematic errors. All measurements were taken using the software ImageJ version 1.52a (Schneider et al. 2012) on size-calibrated photographs.

Our morphometric analysis consisted of multiple analysis steps. First, we performed a reliability analysis. For this purpose, we aligned and photographed 16 characters in a total of 17 females twice and then calculated measure reliability (which is 1-measurement error, see László et al. 2013) (detailed in Table S4). All characters with reliability below 85% were discarded (one in total). Measurement values of the retained 15 characters in all 223 females samples were studied in a multivariate ratio analysis (MRA) as outlined by Baur and Leuenberger (2011, 2020). We computed an isometric size axis (isosize) and then performed a shape PCA in order to determine whether or not a morphometric pattern corresponds to groups. We also checked measurements for exhibiting allometry by plotting isosize against shape PCs. All statistical analyses were conducted with the software R version 3.3.3 (R Core Team 2017) and using custom R scripts provided by Baur and Leuenberger (2020). Missing values were replaced by using the function “mice”, which applies chained equations, of the R package “mice” and using the function’s default options (van Buuren and Groothuis-Oudshoorn 2011). More details on the microscopic photographs, the obtained measurements, and the applied workflow used for imaging and measuring are given in the Supplementary Tables S5.

## Results

### Cuticular hydrocarbons of dry-mounted and of fresh samples

In 947 of the 1,042 studied dry-mounted wasps (91%) was the amount of extracted CHCs sufficient to identify the wasp’s chemotype. Note that our sampling included also 357 males, which consistently showed only one chemotype (similar to the females’ chemotype 1). The CHC extracts of 42 males proofed to be insufficient for characterizing their CHC profile. Some of the wasps whose chemotype we were able to determine had been collected more than 200 years ago (the oldest were collected in 1826), indicating that even old dry-mounted samples can represent a valuable resource in the field of chemical ecology. We found a statistically significant correlation between the age of the dry-mounted samples and the ability to determine their chemotype (logistic regression, *p* value = 6.14e-06, Fig. S3). However, not all recently collected dry-mounted samples contained a sufficient amount of CHCs to characterize their CHC profile and to determine their chemotype, suggesting that additional factors impact CHC preservation. In contrast, most of the CHC samples collected or probed specifically for CHC analyses (311 out of 324 in total, 13 of which were of insufficient quality, probably because the cold chain was interrupted during sample shipment; these samples were hence discarded by us) were suitable to identify the wasps’ chemotype.

Among the 311 samples that were freeze-killed or probed with SPME fibers, there was a clear separation between the two chemotypes based on the exclusive presence of *udp* alkenes in chemotype 1 females and the almost exclusive presence of *edp* alkenes in chemotype 2 females (*udp* alkenes were only found in traces, if at all). The CHC profile of dry-mounted samples belonging to chemotype 1 (694) also exclusively showed *udp* alkenes. However, almost all of the remaining dry-mounted females (245), here considered expressing chemotype 2, showed a mix of *edp* and *udp* alkenes. In fact, only seven (three collected in 2012 and four in 2017, respectively, by M. Halada and by Z. Haladova) showed CHC profiles identical to those of freeze-killed females expressing chemotype 2. Applying a derivatization protocol to CHC extracts from a subset of 92 dry-mounted females here considered having expressed chemotype 2, we found a mix of *edp* and *udp* alkenes in the CHC profiles of 86 samples (Table S3). Specifically, we found this mix, depending on the sample, at hydrocarbon chain positions 23 and 25 and, less frequently, at position 27. However, we never found *udp* alkenes at hydrocarbon chain position 29 (Fig. S4, Table S3). In only six of the 92 dry-mounted samples studied in detail did we find exclusively *edp* alkenes — thus a CHC profile composition identical to that obtained from freeze-killed fresh samples expressing chemotype 2. Since the CHC extracts of most dry-mounted females not having expressed chemotype 1 contained a mix of *udp* and *edp* alkenes and because we never found the deviant CHC profile having been expressed by freshly collected females, sampled at locations spatially close to those from which we had dry-mounded samples with a deviant chemotype 2 profile, we assumed that the deviation is an artifact.

### Spatial Differences in Chemotype Frequencies

We found females of both chemotypes in the Western Palearctic and in the Eastern Palearctic, the presumed glacial refuge of *O. spinipes*, which shows the typical distribution of a Euro-Siberian faunal element as defined by de Lattin (1967). However, we discovered significant geographic structure in the frequency of the two chemotypes, with one chemotype being more frequent in some regions than the other (Fig. 2). For example, we recorded only females of chemotype 1 in western France, the UK, Italy, and the southern parts of Switzerland. In Central Europe, we found a conspicuous North-South gradient in the two chemotype frequencies, with females expressing chemotype 2 being more frequent in the North than females expressing chemotype 1 and vice versa (shift at a latitude of ∼ 49° N). In the South-West of Austria, almost all females express chemotype 1, while in Eastern Austria both chemotypes occur at about the same frequency. In the Czech Republic, we found most females to express chemotype 2, except in regions spatially close to Austria and Poland. Here, females expressing chemotype 1 were more frequent. Interestingly, although the majority of females from northern Central Europe expressed chemotype 2, this chemotype is almost absent at least in parts of Northern Europe (Estonia and Sweden). We found exclusively females expressing chemotype 1 in Kazakhstan, Mongolia, and parts of Russia (Siberia and Ural). While females expressing chemotype 2 seem to be absent (or at least rare) in Central Russia, we found most females expressing chemotype 2 at Russia’s Pacific coast, as well as in Eastern Turkey and in Romania. The two females in our dataset collected in Bulgaria also expressed chemotype 2.

**Fig. 2.**
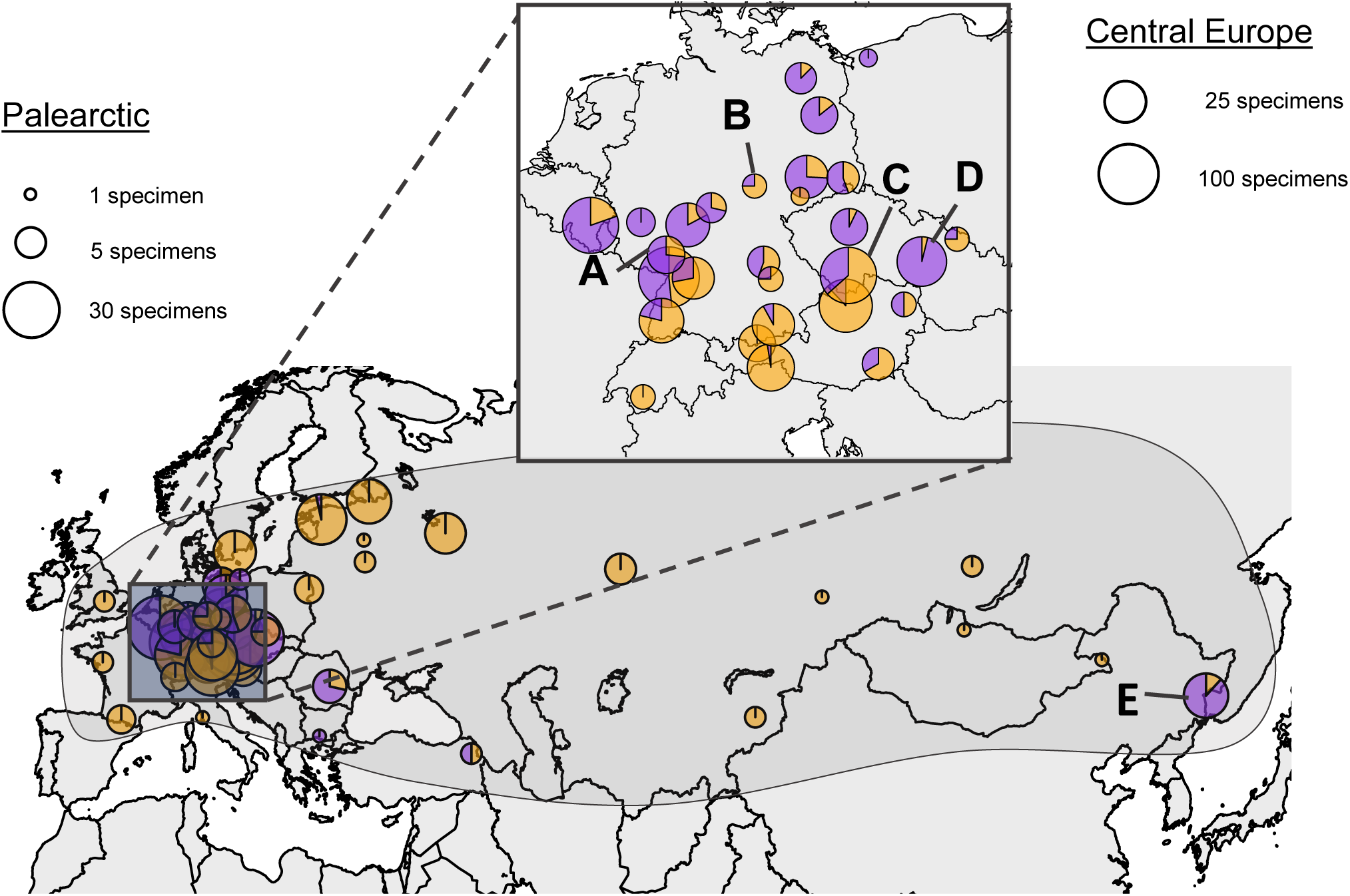
Geographical differences in the frequency at which *Odynerus spinipes* females express chemotype 1 (orange) and chemotype 2 (purple). The diameter of the pie charts is proportional to the number of females analyzed a given location. Note that the pie charts on the two maps have different scales. The approximate distribution of the species is indicated by the continuous black line.

### Morphometric Analysis

The conducted reliability tests indicated that only 15 of the 16 evaluated characters were measured with consistent accuracy (Table 1). Multivariate ratio analysis (MRA) of the 15 characters in 82 females of chemotype 1 and 141 females of chemotype 2 revealed no statistically significant differences between females expressing chemotype 1 and females expressing chemotype 2 (including those with deviant CHC profile) in the four selected populations, neither in size nor in shape axes: Fig. 3 shows that groups are largely overlapping. However, we observed that *O. spinipes* females from the Far East of Russia (locality E) are on average slightly larger than females collected in Europe (localities A– D).

**Fig. 3.**
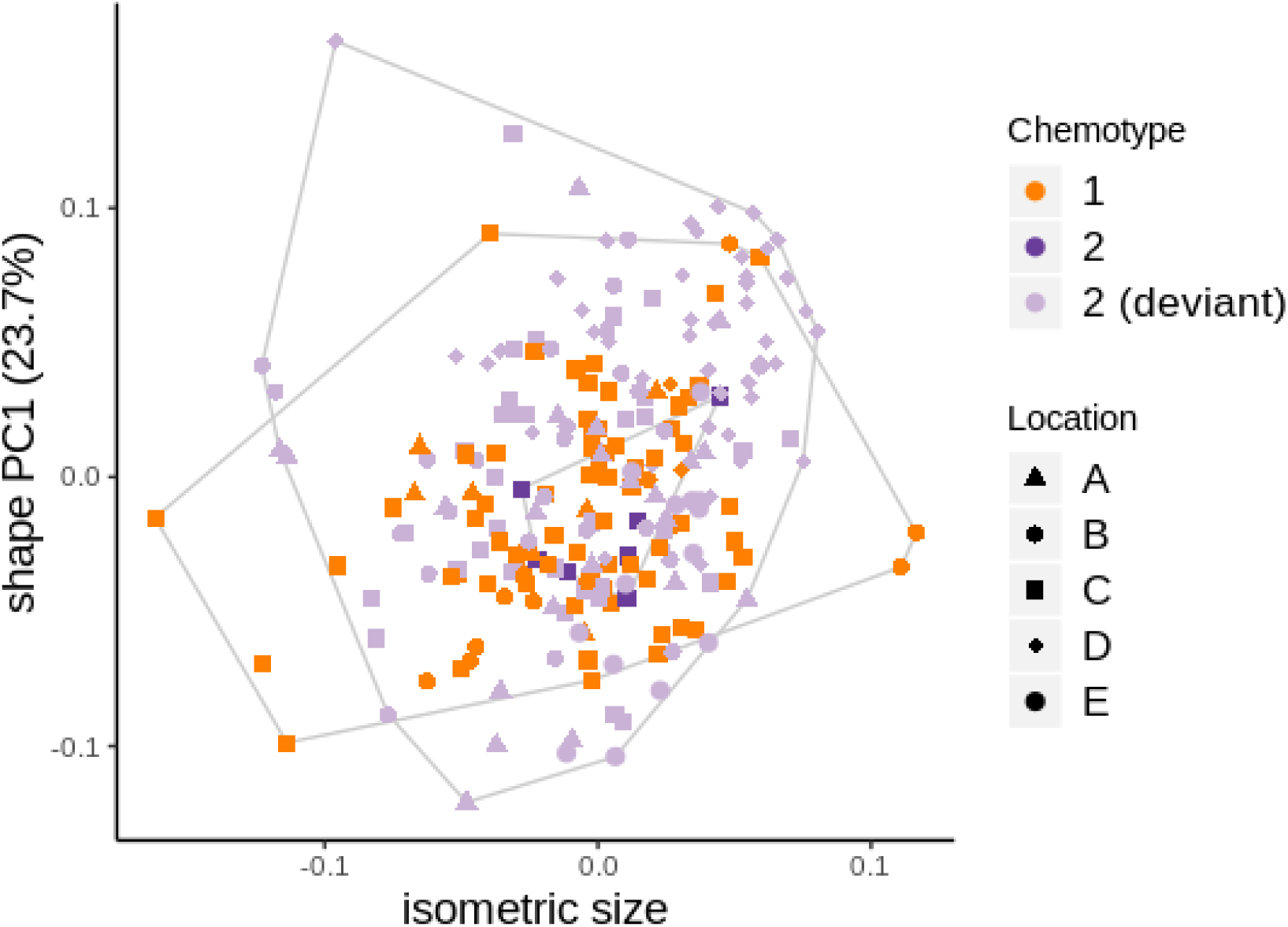
Score plot from a multivariate ratio analysis (MRA) of distance measurements showing first shape principal component versus isometric size. Color: orange, chemotype 1, purple, chemotype 2, light purple, deviant chemotype 2. Shapes represent each a locality: Frankfurt a. M. (A, triangles), Leipzig (B, circles), Southern Bohemia (C, squares), Syrovice (D, diamonds), Far East (E, squares with a cross).

## Discussion

### Temporal Chemical Stability of Cuticular Hydrocarbons

The results of our study highlight the potential value of dry-mounted insects stored in private and public collections for chemical ecological studies. Our data demonstrate the long-term chemical stability of CHCs on specimens. Long-term stability of CHCs had previously been reported by Page et al. (1990) and by Martin et al. (2009b) who analyzed samples that were up to 70- and 20-years-old, respectively. Since CHCs are typically species-specific, their analysis on museum specimens opens for the door for inferring the identity of samples (including type material) that otherwise could perhaps only be identified using DNA barcoding or that could not be identified at all. The utility of CHC for identifying cryptic species has been shown in numerous studies (Haverty et al. 1990; Page et al. 1997; Akino et al. 2002; Lucas et al. 2002; Schlick-Steiner et al. 2006; Martin et al. 2008; Guillem et al. 2012; Vaníčková et al. 2014). And while the ability to study CHCs in dry-mounted insects has been demonstrated in earlier studies (Bartelt et al. 1986; Jallon and David 1987; Grunshawn et al. 1990; Page et al. 1990; Chapman et al. 1995; Everaerts et al. 1997; Clément et al. 2001; Dapporto et al. 2004; Symonds and Elgar 2004; Uva et al. 2004; Dapporto 2007; Hay-Roe et al. 2007; Martin et al. 2008; Martin et al. 2009a, b), these studies relied on samples that had been comparatively recently collected (except that published by Page et al. 1990).

The analysis of CHC extracts of dry-mounted specimens has limitations. First, CHC extracts from such samples have a high chance of being contaminated. While most of the contaminations will likely be volatile non-CHCs, it is imaginable that CHCs from other samples stored in the same box get transferred over the course of time on the cuticle of samples of interest. Unless fresh material is unavailable, dry-mounted samples should therefore be avoided for *de novo* characterizing the CHC profile of a species. If such samples are nonetheless used, the obtained chromatograms should be interpreted with great caution. A second limitation of dry-mounted samples for chemical analyses is the low abundance at which the CHCs are obtained. Since we found in initial tests the trend for longer extraction duration time to increase the yield of CHCs, we recommend future studies on the CHC of dry-mounted to extract CHC for at least for 10 minutes. One likely reason for the low amount of CHCs on dry-mounted samples is evaporation of the insect’s CHCs over time, or that the hydrophobic compounds diffuse into the now dried interior of the specimen. The fact that we successfully extracted CHC also from very old (> 100 years) samples suggests that the effect of evaporation is likely comparatively small, or at least to some extent predictable. A potentially more severe factor than specimen age is the kind of chemical used to kill and/or preserve the insects, as some of these chemicals are powerful organic solvents (e.g., carbon tetrachloride, ethanol, diethyl ether, ethyl acetate; Gibb and Oseto 2019). If the insects were killed by being immerged in organic solvent, their CHCs are likely removed very efficiently from the cuticle, since most hydrocarbons are soluble in organic solvents (Krogmann and Holstein 2010; Tewari and Vishnoi 2017). Even if not submerged in solvent, collecting several specimens in the same vial containing an organic solvent as killing agent could potentially lead to mixing of CHC’s between specimens. By contrast, freeze-killing of insects is generally assumed to have no notable impact the insects’ CHCs. We attribute the chemicals used to kill the insects studied by us as the most likely cause of why we were in some instances unable to extract notable quantities of CHCs regardless of how long ago a sample had been collected.

The most severe problem encountered when studying CHC extracts from old dry-mounted specimens is that the CHC profile of samples can be slightly altered over time. Specifically, we found the CHC profiles of various *O. spinipes* females to contain a mix of *udp* and *edp* alkenes (same chain length; 23, 25, 27). In contrast, the CHC profiles of freeze-killed specimens consistently contained either *udp* alkenes or *edp* alkenes. In some females of chemotype 2, we found traces of *udp* alkenes, but these alkenes were never as abundant as *edp* alkenes of the same chain length (see also results presented by Wurdack et al. 2015). We exclude the possibility that samples with deviant CHC profile represent a cryptic species based on the results from the morphometric analyses. We also consider it extremely unlikely that these deviant CHC profiles represent a third (and intermediate) chemotype, because we found this chemotype exclusively in CHC extracts originating from dry-mounded samples, some of which collected at locations close to sites where we freshly collected wasps without ever encountering such a deviant profile. It therefore appears more plausible to us that the CHC profile of a significant portion of the dry-mounted wasps had changed. It is known that the volatility of alkenes is generally higher than that of alkanes (including methyl-branches ones) and that the volatility of alkenes differs from each other depending on the molecule’s length and the position of the double bond (Gibbs and Pomonis 1995; Tewari and Vishnoi 2017). A difference in volatility between alkanes and alkenes could explain why we always observed alkanes in the studied dry-mounted specimens. Additionally, it is known that alkene double bonds can alter their position by rearrangement reactions and that such reactions can be triggered by light (Tewari and Vishnoi 2017), which might have happened in insect boxes where the dry-mounted samples were stored. Finally, it is also possible that the *udp* alkenes found in the deviant chemotype 2 came from sources inside the dry-mounted samples. Because of their dryness, it is possible that hexane extracts CHCs also inside the specimens and not only on the cuticle of the specimens as it does when processing freeze-killed fresh samples. Incidentally, Martin et al. (2009b) also reported a reduction in the quantity of an alkene (c31ene) in CHC extracts from hornets collected 20 years ago compared to CHC extracts from freshly-collected hornets (*Vespa* spp.).

### Amount and Spatial Distribution of Different CHC Chemotypes

Based on samples from three field sites in southern Germany, Wurdack et al. (2015) reported *O. spinipes* females to be able to express two different chemotypes and conspecific males to only express one chemotype. The results of our study are fully consistent these observations, but our conclusions are based on a significantly larger sample size and covers samples collected from a significantly larger fraction of the species’ geographical range. However, in contrast to the observations made by Wurdack et al. (2015), we found the frequency of the two chemotypes expressed by females to geographically vary considerably and to be rarely balanced. Most notably, we found females expressing both chemotypes to also occur in the Eastern Palearctic at the Pacific coast — the presumed glacial refuge of this Euro-Siberian faunal element.

The chemotype frequency differences in some parts of the distributional range of *O. spinipes* are remarkable, and multiple mechanisms are imaginable that could have given rise to them. Yet, because species population structures are the result of both present and historical processes (Hewitt 1999), it is difficult to disentangle what factors shaped the chemotype frequency differences in different populations and geographic regions in the absence of additional data (e.g., population genetic information). For example, chemotype frequency differences could simply be the results of genetic drift during range expansion from a single glacial refuge in the Eastern Palearctic. Chemotype frequency differences could additionally reflect different routes and times of the colonization of the Western Palearctic. Finally, chemotype frequency differences could be the result of historical differences in the selection pressure for a given chemotype. The latter aspect if particular intriguing, given that the two chemotypes are chemically mimicked by two cuckoo wasp species (i.e., *Chrysis mediata* and *Pseudochrysis neglecta*), which occur across most (if not all) of *O. spinipes*’ distributional range (Kimsey and Bohart 1991; Yildirim and Strumia 2006; Kurzenko and Lelej 2007; Rosa et al. 2014; Belokobylskij and Lelej 2017; Rosa 2019). Population genetic studies on *O. spinipes* and its kleptoparasitic cuckoo wasps could help disentangling the above factors for having shaped geographic chemotype frequency differences. The presented data provide important information where to geographically preferentially sample *O. spinipes* for such population genetic analyses. They also set the base for investigations that aim at understanding the genetics of the astounding qualitative CHC dimorphism that *O. spinipes* females exhibit.

## Acknowledgments

We are indebted to Fritz Gusenleitner, Rolf Franke, Lukas Kirschey, Lars Krogmann, Andrew Liston, Mikhail Mokrousov, Michael Ohl, Patricia Peters, Karl-Heinz Schmalz, Christian Schmid-Egger, Stefan Schmidt, Martin Schwarz, and Dominique Zimmermann for providing us samples and granting permission to chemically analyze them. We acknowledge Paolo Rosa and Manuela Sann for bringing us in contact with some entomologists and Jean-Yves Baugnée for providing information where to collect *O. spinipes* in Belgium. We are thankful to Rainer Blum for help with georeferencing samples and to Wolf Haberer for help using the GC-MS. ON and VM acknowledge the Struktur-und Genehmigungsbehörde Süd and the Struktur-und Genehmigungsdirektion Nord (both Rhineland Palatinate) for granting permission to collect samples. Part of the present study were funded by the German Research Foundation (DFG9 NI1387/2-1, SCHM 2645/6-1). This work was in part supported by the Russian Foundation for Basic Research (project No. 19–04–00027) and Russian State Research Project No. АААА–А19– 119020690101–6 for SAB.

## References

Akino T, Terayama M, Wakamura S, Yamaoka R (2002) Intraspecific variation of cuticular hydrocarbon composition in *Formica japonica* Motschoulsky (Hymenoptera: Formicidae). Zool Sci 19:1155–1165

Blomquist CJ, Bagnères AG (2010) Insect hydrocarbons. Biology, biochemistry, and chemical ecology. Cambridge University Press, Cambridge

Bagnères AG, Wicker-Thomas C (2010) Chemical taxonomy with hydrocarbons. In: Blomquist GJ, Bagnères AG (eds) Insect hydrocarbons. Biology, biochemistry, and chemical ecology. Cambridge University Press, Cambridge, pp 121–162

Bartelt RJ, Armold MT, Schaner AM, Jackson LL (1986) Comparative analysis of cuticular hydrocarbons in the Drosophila virilis species group. Comp Biochem Phys B 83:731–742

Baur H, Leuenberger C (2011) Analysis of ratios in multivariate morphometry. Sys Biol 60:813–825

Baur H, Leuenberger C (2020) Multivariate Ratio Analysis (MRA): R-scripts and tutorials for calculating Shape PCA, Ratio Spectra and LDA Ratio Extractor (Version 1.02). Zenodo, http://doi.org/10.5281/zenodo.3892267

Bechshøft TØ, Rigét FF, Wiig Ø, Sonne C (2008) Fluctuating asymmetry in metric traits; a practical example of calculating asymmetry, measurement error, and repeatability. Ann Zool Fenn 45:32–38

Belokobylskij SA, Lelej AS (2017) Annotated Catalogue of the Hymenoptera of Russia, Volume I, Symphyta and Apocrita: Aculeata. Proceedings ZIN, 321:1–475

Carlson DA, Roan CS, Yost RA, Hector J (1989) Dimethyl disulfide derivatives of long chain alkenes, alkadienes, and alkatrienes for gas chromatography/mass spectrometry. Anal Chem 61:1564–1571

Chapman RF, Espelie KE, Sword GA (1995) Use of cuticular lipids in grasshopper taxonomy: A study of variation in *Schistocerca shoshone* (Thomas). Biochem Syst Ecol 23:383–398

Clément JL, Bagnères AG, Uva P, Wilfert L, Quintana A, Reinhard J, Dronnet S (2001) Biosystematics of *Reticulitermes* termites in Europe: morphological, chemical and molecular data. Insectes Soc 48:202–215

Cuvillier-Hot V, Cobb M, Malosse C, Peeters C (2001) Sex, age and ovarian activity affect cuticular hydrocarbons in *Diacamma ceylonense*, a queenless ant. J Insect Physiol 47:485–493

Dunkelblum E, Tan TS, Silk PJ (1985) Double-bond location in monounsaturated fatty-acids by dimethyl disulfide derivatization and mass-spectrometry—application to analysis of fatty-acids in pheromone glands of four Lepidoptera. J Chem Ecol 11:265–277

De Oliveira CC, Manfrin MH, de M Sene F, Jackson LL, Etges WJ (2011) Variations on a theme: diversification of cuticular hydrocarbons in a clade of cactophilic *Drosophila*. BMC Evol Biol 11:179

Dapporto L, Palagi E, Turillazzi S (2004) Cuticular hydrocarbons of *Polistes dominulus* as a biogeographic tool: a study of populations from the Tuscan Archipelago and surrounding areas. J Chem Ecol 30:2139–2151

Dapporto L (2007) Cuticular lipid diversification in *Lasiommata megera* and *Lasiommata paramegaera*: the influence of species, sex, and population (Lepidoptera: Nymphalidae). Biol J Linn Soc 91:703–710

de Lattin G (1967) Grundriss der Zoogeographie. G. Fischer (ed), Stuttgart

Dobson AJ (1990) An Introduction to Generalized Linear Models. London: Chapman and Hall

Everaerts CL, Farine JP, Brossut R (1997) Changes of species specific cuticular hydrocarbon profiles in the cockroaches *Nauphoeta cinerea* and *Leucophaea maderae* reared in heterospecific groups. Entomol Exp Appl 85:145–150

Gerritsen H (2013) mapplots: data visualisation on maps. R package version 1.4. http://CRAN.R-project.org/package=mapplots

Gibb TJ, Oseto C (2019) Insect Collection and Identification: Techniques for the Field and Laboratory. Academic Press

Gibbs AG, Pomonis JG (1995) Physical properties of insect cuticular hydrocarbons: The effects of chain length, methyl-branching and unsaturation. Comp Biochem Phys A, 112B:243–249

Gibson GAP (1997) Morphology and Terminology. Annotated Keys to the Genera of Nearctic Chalcidoidea (Hymenoptera) (ed. by Gibson GAP, Huber JT, Woolley JB), pp 16–44. NRC Research Press, Ottawa

Greene MJ, Gordon DM (2003) Cuticular hydrocarbons inform task decisions. Nature 423:32–32

Grunshawn JP, Guermouche H, Guermouche S, Jago ND, Jullien R, Knowles E, Perez F (1990) Chemical taxonomic studies of cuticular hydrocarbons in locusts of the *Schistocerca americana* complex: chemical relationships between New World and Old World species. J Chem Ecol 16:2835–3858

Guillem RM, Drijfhout FP, Martin SJ (2012) Using chemo-taxonomy of host ants to help conserve the large blue butterfly. Biol Conserv 148:39–43

Gusenleitner J (1998) Bestimmungstabellen mittel-und südeuropäischer Eumeniden (Vespoidea, Hymenoptera) Teil 8. Die Gattungen *Odynerus* LATREILLE 1802, *Gymnomerus* BLÜTHGEN 1938, *Paragymnomerus* BLÜTHGEN 1938 und *Tropidodynerus* BLÜTHGEN 1939, Linzer biol Beitr:163–181

Haverty MI, Nelson LJ, Page M (1990) Cuticular hydrocarbons of four populations of *Coptotermes formosanus* Shiraki in the United States. J Chem Ecol 16:1635–1647

Hay-Roe MM, Lamas G, Nation JL (2007) Pre-and postzygotic isolation and Haldane rule effects in reciprocal crosses of *Danaus erippus* and *Danaus plexippus* (Lepidoptera: Danainae), supported by differentiation of cuticular hydrocarbons, establish their status as separate species. Biol J Linn Soc 91:445–453

Hewitt GM (1999) Post-glacial re-colonization of European biota. Biol J Linn Soc 68:87–112

Howard RW, Blomquist GJ (2005) Ecological, behavioral, and biochemical aspects of insect hydrocarbons. Ann Rev Entomol 50:371–393

Hugo LE, Kay BH, Eaglesham GK, Holling N, Ryan PA (2006) Investigation of cuticular hydrocarbons for determining the age and survivorship of Australasian mosquitoes. Am J Trop Med Hyg 74:462–474

Ichinose K, Lenoir A (2009) Ontogeny of hydrocarbon profiles in the ant *Aphaenogaster senilis* and effects of social isolation. C R Biol 332:697–703

Jackson DE, Martin SJ, Ratnieks FL, Holcombe M (2007) Spatial and temporal variation in pheromone composition of ant foraging trails. Behav Ecol 18:444–450

Jallon JM, David JR (1987) Variation in the cuticular hydrocarbons among the eight species of *Drosophila melanogaster* subgroup. Evolution 41:294–302

Kimsey LS, Bohart RM (1991) The Chrysidid Wasps of the World. Oxford Press, New York, 652 pp

Korb J (2018) Chemical fertility signaling in termites: idiosyncrasies and commonalities in comparison with ants. J Chem Ecol 44:818–826

Krogmann L, Holstein J (2010) Preserving and Specimen Handling. In: Eymann J, Degreef J, Häuser CL, Monje JC, Samyn Y, VandenSpiegel D (eds) Manual on field recording techniques and protocols for All Taxa Biodiversity Inventories (ATBIs), part 2, pp 463–481

Kuo TH, Yew JY, Fedina TY, Dreisewerd K, Dierick HA, Pletcher SD (2012) Aging modulates cuticular hydrocarbons and sexual attractiveness in *Drosophila melanogaster*. J Exp Biol 215:814–821

Kurzenko NV, Lelej AS (2007) Fam. Chrysididae-Chrysidid wasps. Key to the Insect of Russian Far East, Vol. 4, Part 5. Dalnauka, Vladivostok, pp 998–1006

László Z, Baur H, Tóthmérész B (2013) Multivariate ratio analysis reveals *Trigonoderus pedicellaris* Thomson (Hymenoptera, Chalcidoidea, Pteromalidae) as a valid species. Syst Entomol 38: 753–762

Linsenmaier W (1951) Die europäischen Chrysididen (Hymenoptera). Versuch einer natürlichen Ordnung mit Diagnosen. Mitteilungen der Schweizerischen Entomologischen Gesellschaft 24:1–110

Lucas C, Fresneau D, Kolmer K, Heinze J, Delabie JH, Pho DB (2002) A multidisciplinary approach to discriminating different taxa in the species complex *Pachycondyla villosa* (Formicidae). Biol J Linn Soc 75:249–259

Marten A, Kaib M, Brandl R (2009) Cuticular hydrocarbon phenotypes do not indicate cryptic species in fungus-growing termites (Isoptera: Macrotermitinae). J Chem Ecol 35:572–579

Martin SJ, Helanterae H, Drijfhout FP (2008) Evolution of species-specific cuticular hydrocarbon patterns in *Formica* ants. Biol J Linn Soc 95:131–140

Martin SJ, Drijfhout FP (2009a) Nestmate and task cues are influenced and encoded differently within ant cuticular hydrocarbon profiles. J Chem Ecol 35:368–374

Martin SJ, Zhong W, Drijfhout FP (2009b) Long-term stability of hornet cuticular hydrocarbons facilitates chemotaxonomy using museum specimens. Biol J Linn Soc 96:732–737

Martin SJ, Carruthers JM, Williams PH, Drijfhout FP (2010) Host specific social parasites (*Psithyrus*) indicate chemical recognition system in bumblebees. J Chem Ecol 36:855–863

Nunes TM, Turatti IC, Lopes NP, Zucchi R (2009) Chemical signals in the stingless bee, *Frieseomelitta varia*, indicate caste, gender, age, and reproductive status. J Chem Ecol 35:1172–1180

Page M, Nelson LJ, Haverty MI, Blomquist GJ (1990) Cuticular hydrocarbons of eight species of North American cone beetles, *Conophthorus* Hopkins. J Chem Ecol 16:1173–1198

Page M, Nelson LJ, Blomquist GJ, Seybold SJ (1997) Cuticular hydrocarbons as chemotaxonomic characters of pine engraver beetles (*Ips* spp.) in the *grandicollis* subgeneric group. J Chem Ecol 23:1053–1099

Palmer A, Strobeck C (1986) Fluctuating asymmetry: measurement, analysis, patterns. Annu Rev Ecol Evol S 17:391–421

Pauli T, Castillo-Cajas RF, Rosa P, Kukowka S, Berg A, van den Berghe E, Fornoff F, Hopfenmüller S, Niehuis M, Peters RS, Staab M, Strumia F, Tischendorf S, Schmitt T, Niehuis O (2019) Phylogenetic analysis of cuckoo wasps (Hymenoptera: Chrysididae) reveals a partially artificial classification at the genus level and a species-rich clade of bee parasitoids. Syst Entomol 44:322–335

Polidori C, Giordani I, Wurdack M, Tormos J, Asís JD, Schmitt T (2017) Post-mating shift towards longer-chain cuticular hydrocarbons drastically reduces female attractiveness to males in a digger wasp. J Insect Physiol 100:119–127

R: A language and environment for statistical computing. R Foundation for Statistical Computing, Vienna, Austria. ISBN 3-900051-07-0, URL http://www.R-project.org/.

Rosa P, Wei NS, Xu ZF (2014) An annotated checklist of the chrysidid wasps (Hymenoptera, Chrysididae) from China. ZooKeys 455:1–128

Rosa P, Pavesi M, Soon V, Niehuis O (2017): *Pseudochrysis* Semenov, 1891 is the valid genus name for a group of cuckoo wasps frequently referred to as *Pseudospinolia* Linsenmaier, 1951 (Hymenoptera, Chrysididae). Deut Entomol Z 64:69–75

Rosa P (2019) New remarkable species in the *Chrysis ignita* group (Hymenoptera, Chrysididae) and an overview on Central Asian species, with new synonymies, Linzer biol Beitr 51:397–417

Schlick-Steiner BC, Steiner FM, Moder K, Seifert B, Sanetra M, Dyreson E, Stauffer C, Erhard C (2006) A multidisciplinary approach reveals cryptic diversity in Western Palearctic *Tetramorium* ants (Hymenoptera: Formicidae). Mol Phylogenet Evol 40:259–273

Schneider C, Rasband W, Eliceiri K (2012) NIH Image to ImageJ: 25 years of image analysis. Nat Methods 9:671–675. https://doi.org/10.1038/nmeth.2089

Shuckard WE (1837) Description of the genera and species of the British Chrysididae. The Entomological Magazine 4:156–177

Sledge MF, Boscaro F, Turillazzi S (2001) Cuticular hydrocarbons and reproductive status in the social wasp *Polistes dominulus*. Behav Ecol Sociobiol 49:401–409

Strohm E, Herzner G, Kaltenpoth M, Boland W, Schreier P, Geiselhardt S, Peschke K, Schmitt T (2008a). The chemistry of the postpharyngeal gland of female European beewolves. J Chem Ecol 34:575–583

Strohm E, Kroiss J, Herzner G, Laurien-Kehnen C, Boland W, Schreier P, Schmitt T (2008b) A cuckoo in wolves’ clothing? Chemical mimicry in a specialized cuckoo wasp of the European beewolf (Hymenoptera, Chrysididae and Crabronidae). Front Zool 5:2

Symonds MR, Elgar MA (2004) The mode of pheromone evolution: evidence from bark beetles. Proc Biol Sci 271:839–846

Tewari KS, Vishnoi NK (2017) A textbook of organic chemistry, 4th edn., Vikas Publishing House.

Uva P, Clément JL, Bagnères AG (2004) Colonial and geographic variations in agonistic behaviour, cuticular hydrocarbons and mtDNA of Italian populations of *Reticulitermes lucifugus* (Isoptera, Rhinotermitidae). Insectes Soc 51:163–170

Van Buuren S, Groothuis-Oudshoorn K (2011) Mice: multivariate imputation by chained equations in RJ Stat. J Stat Softw 45:1–67

Vaníčková L, Virgilio M, Tomcala A, Brízová R, Ekesi S, Hoskovec M, Kalinová B, Do Nascimento RR, De Meyeret M (2014) Resolution of three cryptic agricultural pests (*Ceratitis fasciventris, C. anonae, C. rosa*, Diptera: Tephritidae) using cuticular hydrocarbon profiling. Bulletin Entomol Res 104:631–638

Woydak H (2006) Abhandlungen aus dem Westfälischen Museum für Naturkunde 68. Jahrgang 2006, Heft 1. In: Tenbergen B (ed) Hymenoptera Aculeata Westfalica Die Faltenwespen von Nordrhein-Westfalen (Hymenoptera, Vespoidea; Vespidae und Eumenidae) (Soziale Papier- und Lehmwespen). Landschaftsverband Westfalen-Lippe, Münster, pp 57–60

Wurdack M, Herbertz S, Dowling D, Kroiss J, Strohm E, Baur H, Niehuis O, Schmitt T (2015) Striking cuticular hydrocarbon dimorphism in the mason wasp *Odynerus spinipes* and its possible evolutionary cause (Hymenoptera: Chrysididae, Vespidae). Proc Biol Sci 282:20151777

Yildirim E, Strumia F (2006) Contribution to the knowledge of Chrysididae fauna of Turkey. Part 3: Chrysidinae (Hymenoptera, Chrysididae). Linzer Biologische Beiträge, 39:973–984

