## Supplemental Figure 1 for "Cuticular Hydrocarbons on Old Museum Specimens of the Spiny Mason Wasp, *Odynerus spinipes*, (Hymenoptera: Vespidae: Eumeninae) Shed Light on the Distribution and on Regional Frequencies of Distinct Chemotypes"

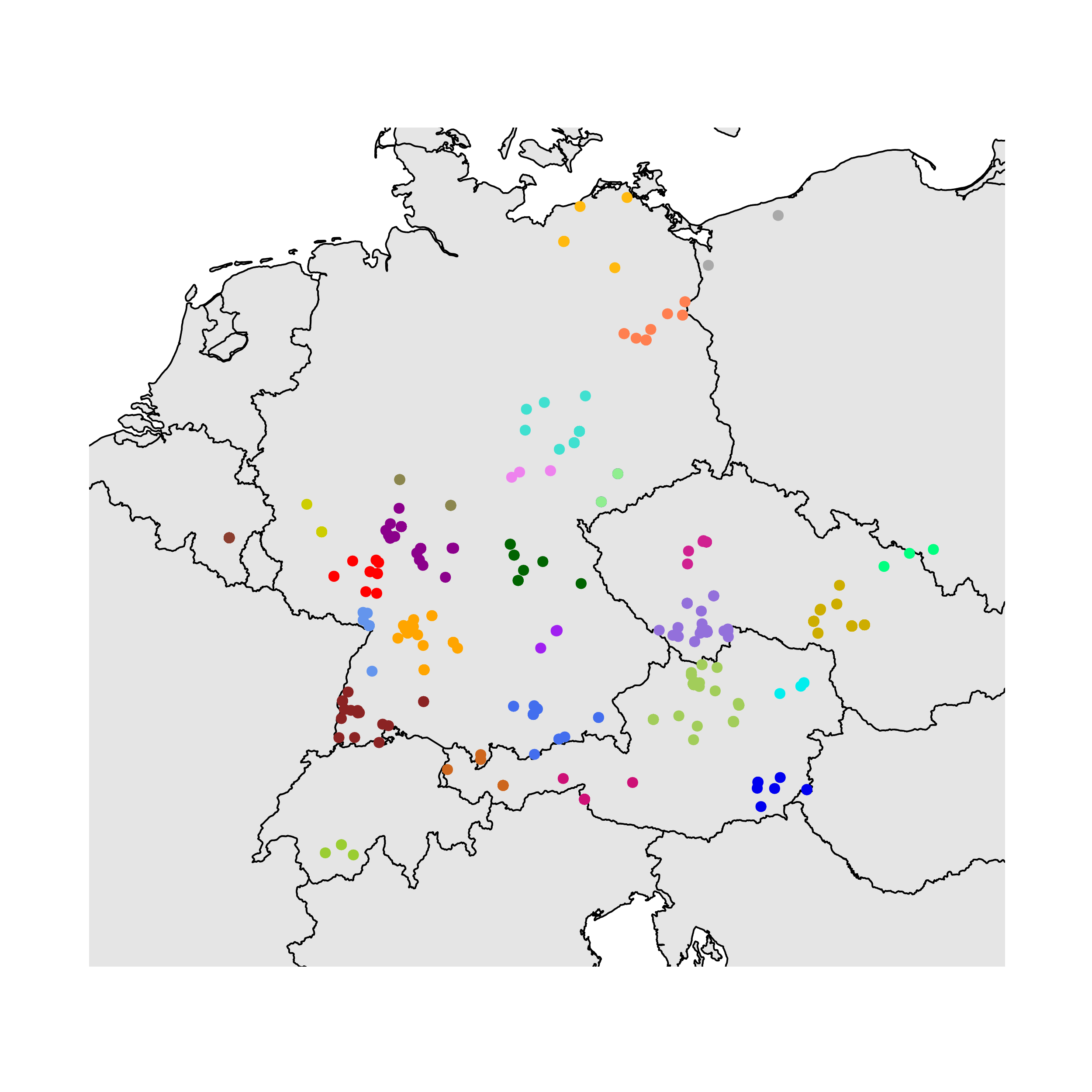

Fig. S1. Map of Central Europe with all localities from which females *Odynerus spinipes* were analyzed. In same color plotted are localities that were grouped together under a standardized area of 12,272 km^2^. Red circles indicate localities from which females were sampled to conduct morphometric analyses.
