## Supplemental Figure 2 for "Cuticular Hydrocarbons on Old Museum Specimens of the Spiny Mason Wasp, *Odynerus spinipes*, (Hymenoptera: Vespidae: Eumeninae) Shed Light on the Distribution and on Regional Frequencies of Distinct Chemotypes"

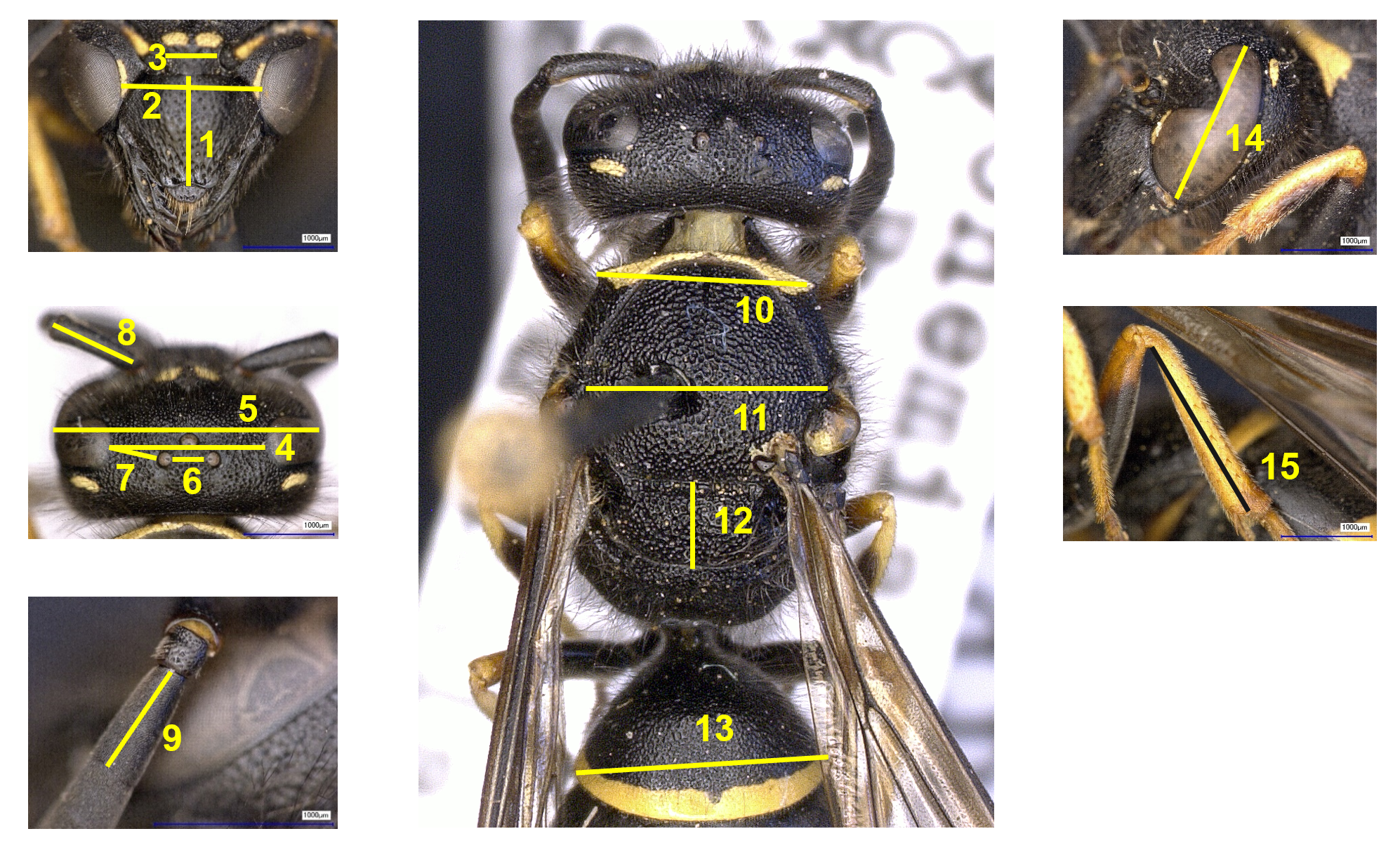

Fig. S2. Measurements used for the morphometric analyses of *Odynerus spinipes* females: clypeus height (1), inner orbit of eye distance (2), toruli distance (3), eye distance (4), head breadth (5), lateral ocelli distance (6), lateral ocellus to eye distance (7), scape length (8), flagellum length (9), pronotal collar breadth (10), mesoscutum breadth (11), scutellum length (12), gastral tergite 1 breadth (13), eye height (14), metatibia length (15). (Photographs by VC Moris and O Niehuis).
