## Supplemental Figure 3 for "Cuticular Hydrocarbons on Old Museum Specimens of the Spiny Mason Wasp, *Odynerus spinipes*, (Hymenoptera: Vespidae: Eumeninae) Shed Light on the Distribution and on Regional Frequencies of Distinct Chemotypes"

Number of samples

Percentage identified (%)

|  | total | percentage |
| --- | --- | --- |
| 1826-1840 | 5 | 100 |
| 1871-1885 | 2 | 100 |
| 1886-1900 | 6 | 83.3333 |
| 1901-1915 | 51 | 88.2353 |
| 1916-1930 | 108 | 74.0741 |
| 1931-1945 | 140 | 87.8571 |
| 1946-1960 | 119 | 98.3193 |
| 1961-1975 | 52 | 86.5385 |
| 1976-1990 | 101 | 92.0792 |
| 1991-2005 | 206 | 93.6893 |
| 2006-2020 | 121 | 97.5207 |

Fig. S3. Number of dry-mounted samples analyzed and percentage of successful chemotype identification according to the years in which the samples were collected.
