## Supplemental Figure 4 for "Cuticular Hydrocarbons on Old Museum Specimens of the Spiny Mason Wasp, *Odynerus spinipes*, (Hymenoptera: Vespidae: Eumeninae) Shed Light on the Distribution and on Regional Frequencies of Distinct Chemotypes"

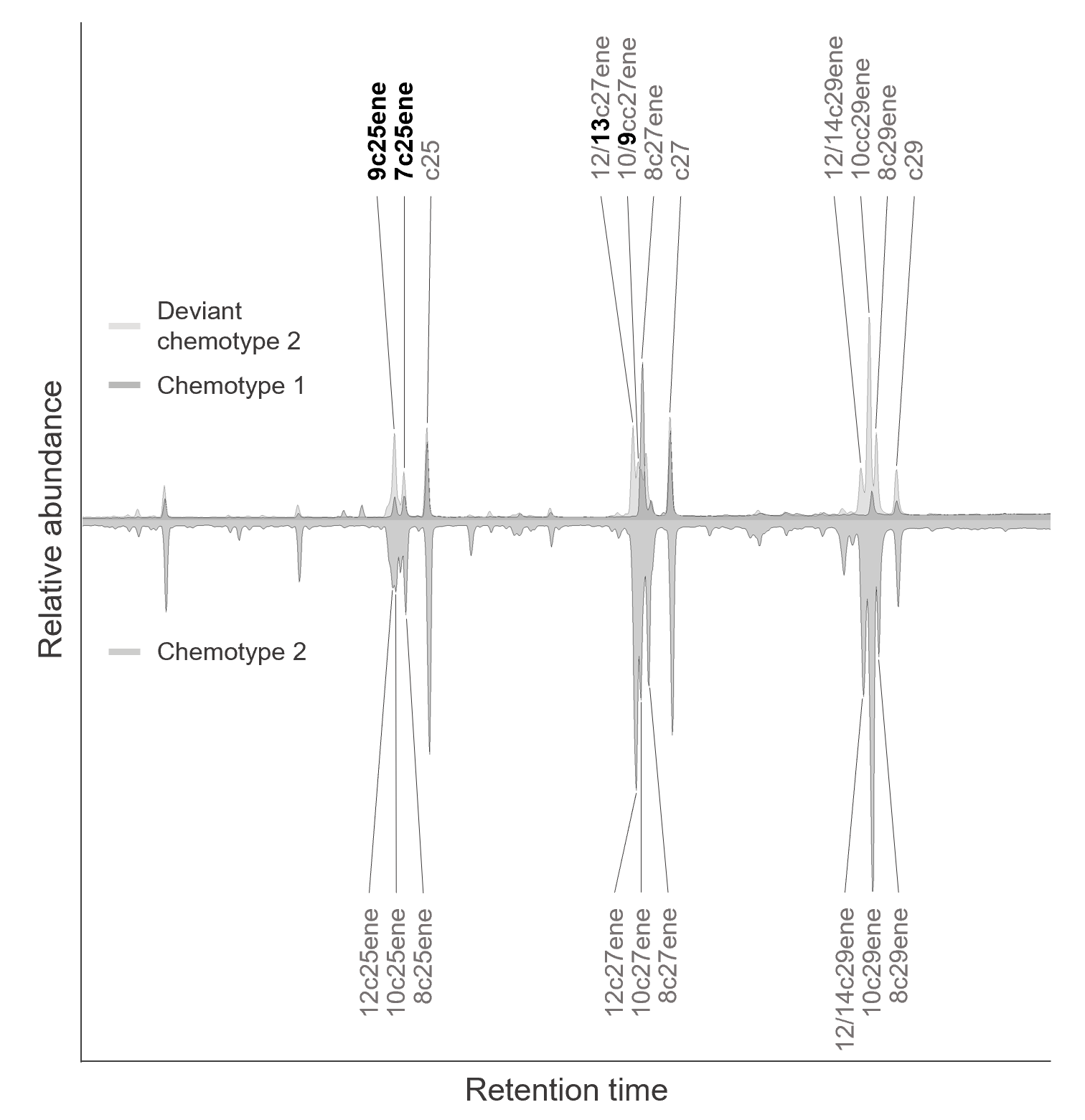

Fig. S4. Chromatograms of CHC extracts from dry-mounted *Odynerus spinipes* females stored in insect boxes. The differences between chemotype 2 and “deviant chemotype 2” are marked in bold: the peaks of c25enes in the “deviant chemotype 2” have a same retention time as those of chemotype 1 (9c25ene and 7c25ene), but there is a clear difference in retention times of c27enes and c29enes in the two CHC profiles.
